# The Murine Neuronal Receptor NgR1 Is Dispensable for Reovirus Pathogenesis

**DOI:** 10.1101/2021.07.22.453368

**Authors:** Pavithra Aravamudhan, Camila Guzman-Cardozo, Kelly Urbanek, Olivia Welsh, Jennifer Konopka-Anstadt, Danica M. Sutherland, Terence S. Dermody

## Abstract

Engagement of host receptors is essential for viruses to enter target cells and initiate infection. Expression patterns of receptors in turn dictate host and tissue tropism and disease pathogenesis during infection. Mammalian orthoreovirus (reovirus) displays serotype-dependent patterns of tropism in the murine central nervous system (CNS) that are dictated by viral attachment protein σ1. However, the receptor that mediates reovirus CNS tropism is unknown. Two proteinaceous receptors have been identified for reovirus, junctional adhesion molecule-A (JAM-A) and Nogo 66 receptor 1 (NgR1). Engagement of JAM-A is required for reovirus hematogenous dissemination but is dispensable for neural spread. To determine whether NgR1 functions in reovirus neuropathogenesis, we compared virus replication and disease following inoculation of wild-type (WT) and NgR1^-/-^ mice. Genetic ablation of NgR1 did not alter replication of neurotropic reovirus strain T3SA- in the intestine and transmission to the brain following peroral inoculation. Viral titers in neural tissues following intramuscular inoculation, which provides access to neural dissemination routes, also were comparable in WT and NgR1^-/-^ mice, suggesting that NgR1 is dispensable for reovirus neural spread to the CNS. The absence of both NgR1 and JAM-A also did not alter replication, neural tropism, and virulence of T3SA- following direct intracranial inoculation. In agreement with these findings, we found that the human but not the murine homolog of NgR1 functions as a receptor and confers efficient reovirus binding and infection of nonsusceptible cells *in vitro*. These results eliminate functions for JAM-A and NgR1 in shaping CNS tropism in mice and suggest that other receptors, yet to be identified, support this function.

**IMPORTANCE:** The CNS presents a range of barriers to pathogen invasion. Yet neurotropic viruses have evolved strategies to breach these barriers and establish infection by engagement of host factors that allow navigation to the CNS and neural cell entry. Human NgR1 was identified as a reovirus receptor in an RNA interference screen and is expressed in CNS neurons in a pattern overlapping with reovirus tropism. Using mice genetically lacking NgR1 expression, and following different routes of inoculation, we discovered that murine NgR1 is dispensable for reovirus dissemination to the CNS, tropism and replication in the brain, and resultant disease. Concordant with these results, expression of human but not murine NgR1 confers reovirus binding and infection of nonsusceptible cells *in vitro*. These results point to species-specific use of alternate receptors by reovirus. A detailed understanding of species- and tissue-specific factors that dictate viral tropism will inform development of interventions and targeted gene delivery and therapeutic viral vectors.

## INTRODUCTION

Binding to host receptors is the first step in viral entry into host cells. While virus-receptor interactions are highly specific, several viruses engage more than a single receptor for cell entry (1,2). The capacity to bind multiple receptors may be required to mobilize the cellular internalization machinery or invade specific tissues in the host. Low-affinity interactions with attachment factors that are abundantly expressed at the cell surface also can promote high-affinity interactions with receptors that mediate cell entry (3,4). Tissue-specific patterns of receptor expression often dictate dissemination routes and tissue tropism and shape disease outcomes during viral infection. Moreover, host-specific receptor expression and polymorphisms can limit infection and transmissibility of viruses between species (5,6). Understanding receptor requirements is essential to identify targets to disrupt virus cell entry and design highly selective viral vectors for therapeutic applications. However, for many viruses, the identity of receptors mediating cell entry and functions of known receptors in pathogenesis remain poorly understood.

Mammalian orthoreovirus (reovirus) displays a broad species tropism and causes serotype-dependent disease in the very young (7). While reovirus infects most humans prior to adolescence (8), severe disease outcomes are rare (9,10). Studies of reovirus infection using mice have established that the routes of transmission following inoculation and tissue tropism in the CNS are serotype-dependent. Following peroral inoculation, primary replication of reovirus occurs in gut-associated lymphoid tissue (11–13). Subsequently, serotype 1 (T1) reovirus spreads through hematogenous routes, infects ependymal cells, and causes nonlethal hydrocephalus (14,15), whereas serotype 3 (T3) reovirus spreads through both hematogenous and neural routes, infects distinct neuronal populations in the CNS, and causes lethal encephalitis (13,15–17). The distinct patterns of CNS tropism displayed by T1 and T3 reovirus are dictated by viral attachment protein σ1 (16,18), suggesting that engagement of specific host receptors by σ1 dictates tropism. While multiple reovirus receptors have been identified, their function in serotype-dependent tropism remains elusive.

Reovirus directly binds sialylated glycans, junctional adhesion molecule-A (JAM-A), and Nogo 66 receptor 1 (NgR1). While reovirus serotypes engage distinct sialylated glycans (19,20), and glycan interactions influence the severity of disease following inoculation, glycan engagement is dispensable for reovirus CNS tropism (21–23). JAM-A is an immunoglobulin superfamily receptor expressed primarily at tight junctions and by hematopoietic cells. Both human and murine homologs of JAM-A are directly bound by the σ1 protein of strains representing all three reovirus serotypes (24,25). JAM-A is required for reovirus to access the blood stream and disseminate hematogenously from the intestine to sites of secondary replication in the host (13). However, if hematogenous dissemination is bypassed by intramuscular or intracranial inoculation, reovirus is capable of transmission through neural routes and can infect the brain of mice lacking JAM-A expression (13). Thus, both sialic acid and JAM-A are dispensable for CNS tropism and neuropathogenesis, suggesting that alternative neural receptors must exist for reovirus.

Human NgR1 was identified as a reovirus receptor in a genome-wide RNA interference screen using HeLa cells (26). NgR1 is a leucine-rich repeat protein primarily expressed on neurons (27). Binding to NgR1 by a variety of structurally dissimilar myelin-associated ligands acts to regulate axonal growth and remodeling (28). The pattern of NgR1 expression in the CNS (29) overlaps with sites of T3 reovirus tropism (13), including neuronal populations in the cerebral cortex, hippocampus, and thalamus. Soluble NgR1 binds reovirus virions, and expression of NgR1 in nonsusceptible Chinese hamster ovary (CHO) cells confers reovirus binding and infection (26). Importantly, infection by T3 reovirus is diminished in primary cultures of neurons derived from cerebral cortices of embryonic NgR1^-/-^ mice (26), suggesting that NgR1 functions as a neural receptor for reovirus. However, the function of NgR1 in reovirus tropism and pathogenesis remained unexplored.

Here, we investigated a function for NgR1 in reovirus neuropathogenesis using mice genetically lacking NgR1 expression. Mice deficient in expression of both JAM-A and NgR1 were used to dissect potential redundant functions for these receptors in reovirus infection. Peroral inoculation was used to mimic the natural fecal-oral route of reovirus transmission, whereas intramuscular or intracranial routes were used to provide direct access to neural routes of dissemination. Regardless of the route of inoculation, we found that NgR1 was dispensable for T3 reovirus spread through both hematogenous and neural routes as well as replication and tropism in the CNS. NgR1 also was not required for T1 reovirus infection of the brain following intracranial inoculation. Concordant with these *in vivo* findings, expression of murine NgR1 did not allow efficient reovirus binding or infection of CHO cells, whereas human NgR1 supported both binding and infection. These findings exclude a function for known murine reovirus receptors in CNS tropism and suggest that alternate receptors must exist to mediate reovirus encephalitis in mice.

## RESULTS

### Murine NgR1 is dispensable for reovirus replication in the intestine and spread to the brain following peroral inoculation

To define the function of NgR1 in reovirus pathogenesis, we tested whether reovirus spread is altered in mice lacking NgR1 expression (NgR1^-/-^) following peroral inoculation, which mimics the natural fecal-oral transmission route of reovirus. The absence of NgR1 expression in knock-out animals was confirmed by immunoblot analysis of brain homogenates from newborn mice (Fig. 1A). Mice deficient in JAM-A alone (JAM-A^-/-^) or both JAM-A and NgR1 (double knock-out [DKO]) were included as controls to exclude the possibility that one receptor might compensate for loss of the other. Neurovirulent reovirus strain T3SA-, which does not bind sialic acid (30), was used in these studies to eliminate the potential confounding variable of sialic acid binding in reovirus pathogenesis (22). Newborn wild-type (WT), NgR1^-/-^, JAM-A^-/-^, and DKO mice were inoculated perorally with 10^4^ plaque forming units (PFU) of reovirus T3SA-. Mice were euthanized at 4-, 8-, and 12-days post-inoculation (dpi), intestine and brain were removed, and titers in homogenates of these tissues were determined using plaque assays. The absence of JAM-A, NgR1, or both receptors did not alter titers in the intestine relative to those in WT mice at any time point examined (Fig. 1B). This result suggests that both JAM-A and NgR1 are dispensable for reovirus replication in the intestine. Titers in the intestine decreased over the experimental time course, likely due to clearance of the virus from this site. Titers in the brain of animals lacking JAM-A were lower than those in WT or NgR1^-/-^ mice (Fig. 1C). These results are consistent with a previous study establishing a requirement for JAM-A in hematogenous reovirus dissemination to the brain following peroral inoculation (13). However, titers in the brain of NgR1^-/-^ mice were comparable to those in WT mouse brain at all time points tested. These results suggest that NgR1 is dispensable for reovirus dissemination to and replication in the brain following peroral inoculation.

**FIG 1.**
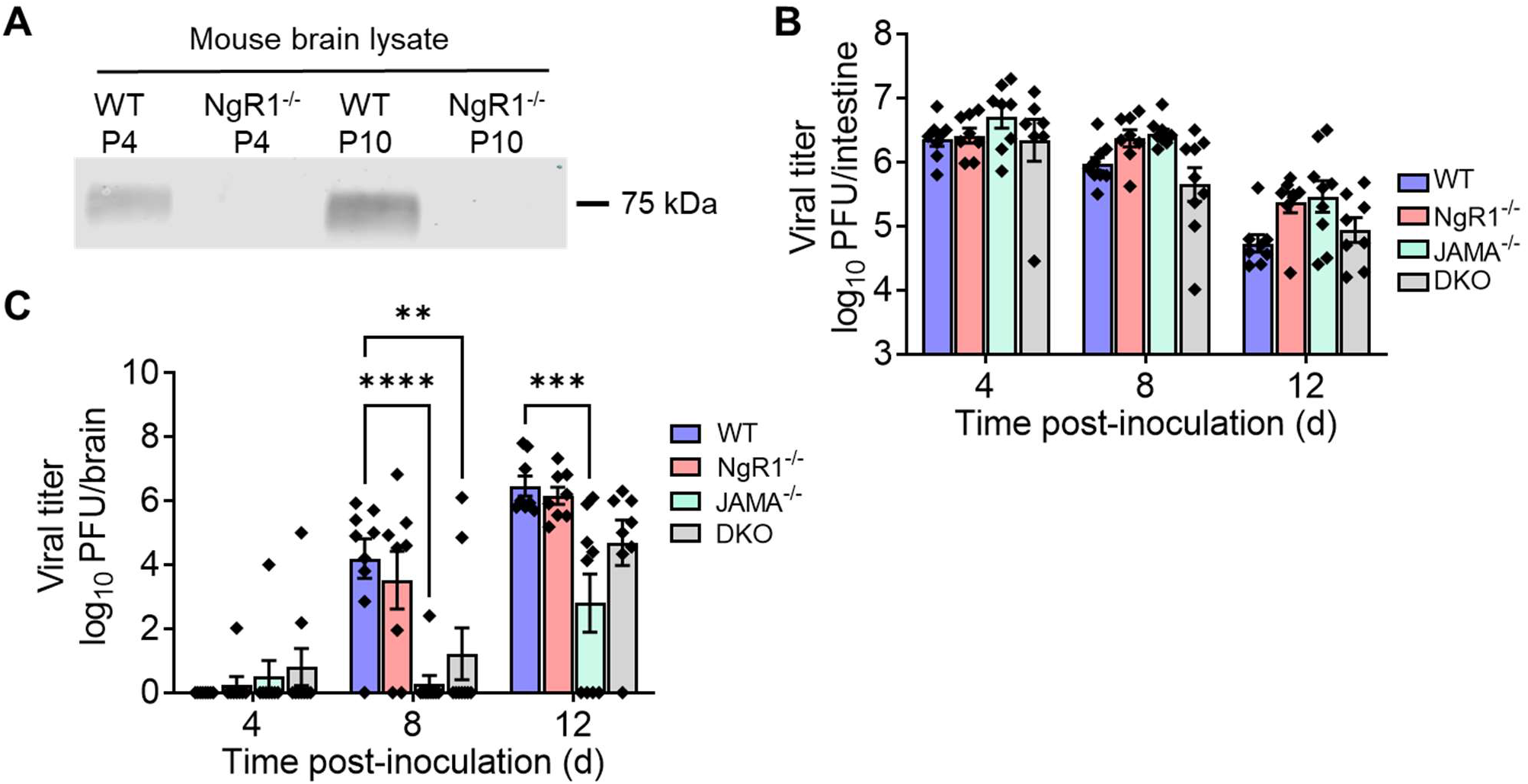
NgR1 is dispensable for reovirus replication and spread to the brain following peroral inoculation. (A) NgR1 expression in brain tissue lysates of WT or NgR1^-/-^ mice at postnatal days 4 (P4) and 10 (P10) were determined by immunoblotting using an antibody specific for mouse NgR1 (AF1440). (B-C) WT, JAM-A^-/-^, NgR1^-/-^, and DKO mice were inoculated perorally with 10^4^ PFU of reovirus T3SA-. Titers in the (B) intestine and (C) brain at the indicated intervals are shown. N = 8 to 9 animals per group for each time point. Error bars indicate SEM. Titers in tissues from knock-out animals were compared with WT at each time point. *, *P* < 0.05; **, *P* < 0.01; ***, *P* < 0.001; ****, *P* < 0.0001; as determined by two-way ANOVA with Dunnett’s multiple comparisons test.

### NgR1 is not required for reovirus replication in the brain and neurovirulence following intracranial inoculation

We next tested whether NgR1 is required for reovirus neurovirulence. For these experiments, we introduced the virus by intracranial inoculation to bypass any function for NgR1 in mediating spread to the brain following inoculation by other routes. Newborn WT, JAM-A^-/-^, NgR1^-/-^, and DKO mice were inoculated intracranially with 25 PFU of T3SA- and monitored for encephalitic symptoms and survival for 21 dpi. Symptoms and severity of neurological disease and survival times of mice lacking NgR1 were comparable to those of WT, JAM-A^-/-^, and DKO animals (Fig. 2A), suggesting that both NgR1 and JAM-A are dispensable for reovirus virulence following intracranial inoculation.

**FIG 2.**
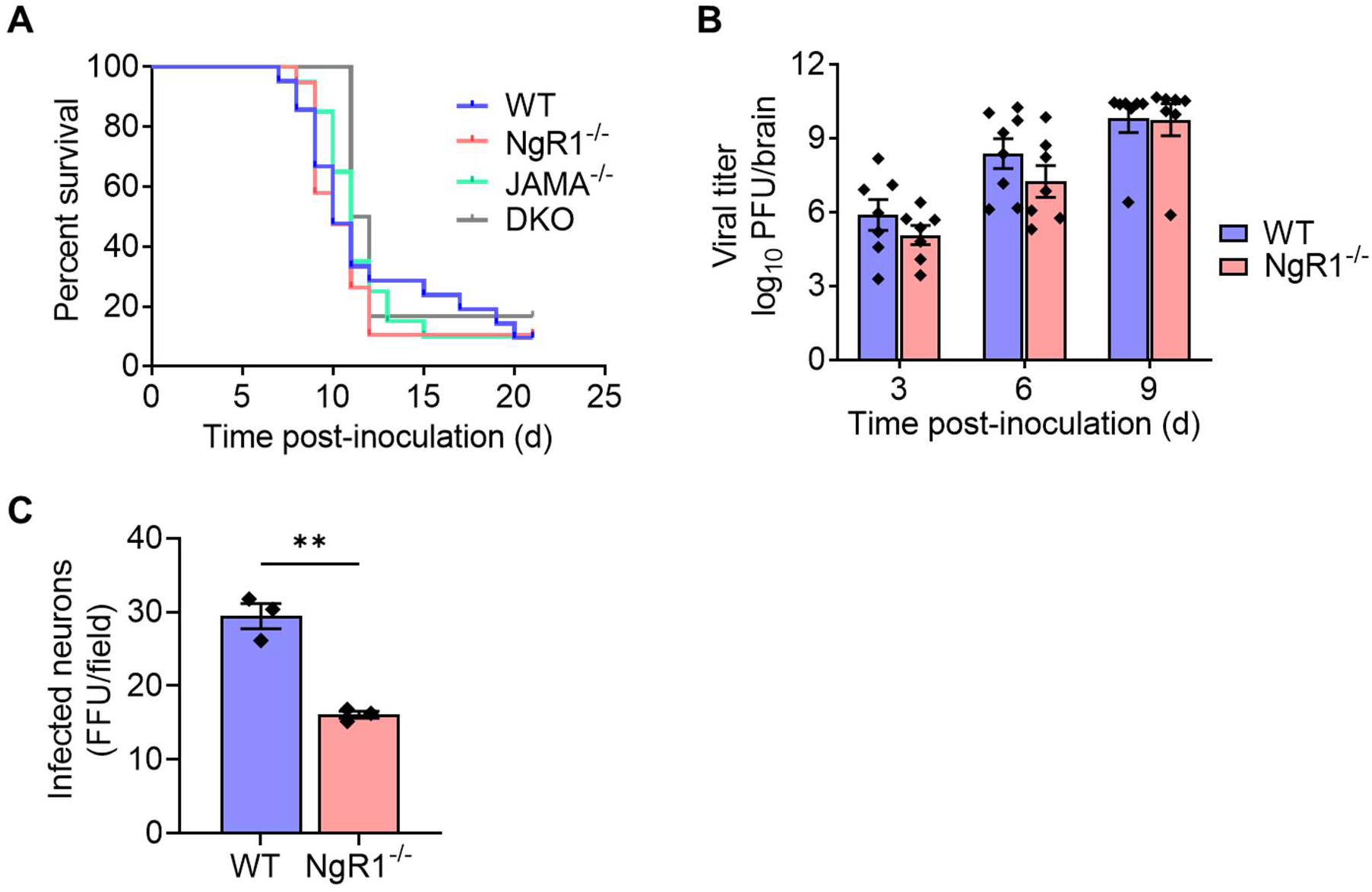
NgR1 is not required for reovirus neuropathogenesis following intracranial inoculation. (A-B) Newborn WT, JAM-A^-/-^, NgR1^-/-^, and DKO mice were inoculated intracranially with 25 PFU of reovirus T3SA-. (A) Mice were monitored for disease signs and survival for 21 d and euthanized when moribund. N = 6 to 21 mice per group. Differences in survival in comparison to WT mice were not significant as determined by log-rank test. (B) At 3, 6, and 9 d after inoculation, mice were euthanized, brains were removed and hemisected, and viral titers in homogenates of the inoculated half of the brain were determined by plaque assay. Results are expressed as mean viral titers. Error bars indicate SEM. N = 8 to 11 mice per group for each time point. Differences in titer between WT and NgR1^-/-^ mice at each time point were not significant (*P* > 0.05) as determined by two-way ANOVA with Dunnett’s multiple comparisons test. (C) Cortical neurons in culture isolated from E15.5 WT or NgR1^-/-^ mice were adsorbed with T3SA+ virions at an MOI of 500 PFU per cell. Cells were fixed at 24 h post-adsorption and immunostained using reovirus-specific antiserum. Infectivity was scored as the mean number of neurons identified based on morphology stained per field-of-view. Bars indicate means and error bars indicate SEM of triplicate samples from one representative experiment out of two conducted. **, *P* < 0.01; determined by *t* test.

Despite the lack of overt differences in disease phenotypes, we hypothesized that the absence of NgR1 might diminish reovirus replication and dissemination in the brain. To test this hypothesis, we inoculated newborn WT and NgR1^-/-^ mice intracranially with 25 PFU of T3SA-. Mice were euthanized at 3, 6, and 9 dpi, brains were removed and hemisected, and titers from the inoculated half of the brain (right hemisphere) were determined. No significant differences in virus titer were observed between WT mice and those lacking NgR1 (Fig. 2B), suggesting that NgR1 is not required for reovirus replication in the brain following intracranial inoculation. These results are in contrast with the significant reduction in reovirus infectivity of cortical neurons cultured from NgR1^-/-^ mice (Fig. 2C) (26).

It is possible that NgR1 mediates spread to and infection of specific neuronal subsets, without affecting overall titers in the brain and neurovirulence. To test this hypothesis, we examined the distribution of reovirus antigen in the brain hemisphere opposing the inoculated hemisphere. Coronal sections of formalin-fixed and paraffin-embedded left-brain hemispheres from the experiments to determine brain viral titers (Fig. 2B) were stained with reovirus-specific antiserum. The intensity of reovirus antigen staining correlated with viral titers from the matched right-brain hemispheres in both WT and NgR1^-/-^ mice (Fig. 3). However, no differences in reovirus antigen staining were observed based on the genotype of the animals. Reovirus-infected neurons, identified based on morphology and antigen staining, were distributed across middle layers of the cortex, CA3 region of the hippocampus, thalamus, and cerebellar Purkinje neurons. The observed antigen distribution is consistent with previous studies of reovirus tropism (13,31) and comparable in WT and NgR1^-/-^ mouse brains. Together, these results suggest that NgR1 is dispensable for reovirus replication, tropism in the brain, and neurovirulence.

**FIG 3.**
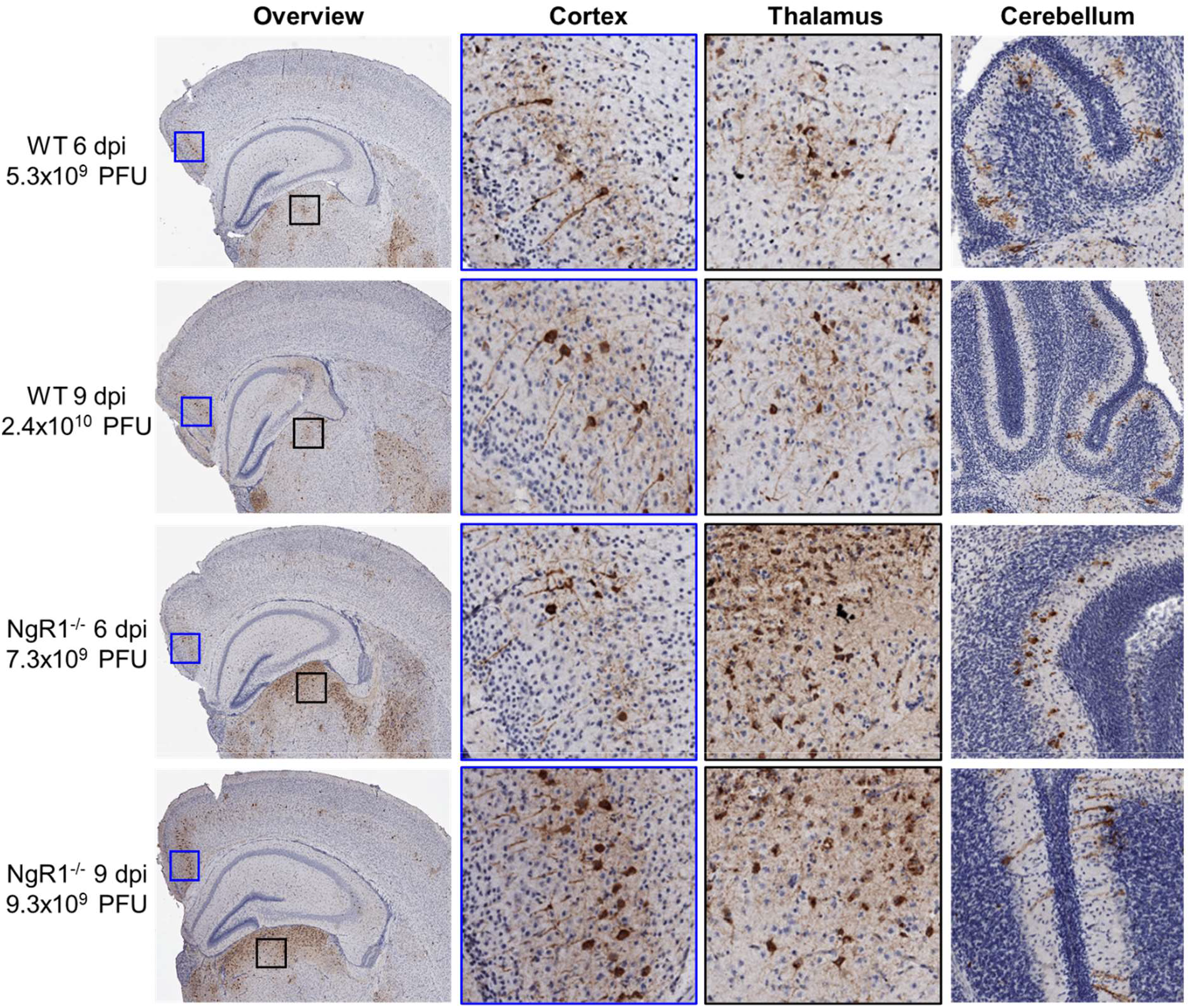
Reovirus tropism in the brain is unaltered in the absence of NgR1. Newborn WT and NgR1^-/-^ mice were inoculated intracranially in the right brain hemisphere with 25 PFU of reovirus T3SA-. Mice were euthanized 6- or 9-d post-inoculation (dpi), and brains were removed and hemisected. Left-brain hemispheres were fixed in formalin and embedded in paraffin. Coronal sections of the left-brain hemisphere were stained with reovirus-specific antiserum and hematoxylin. Representative sections show reovirus antigen in cortex, hippocampus, thalamus, and cerebellum. Enlarged images of areas boxed in the overview show reovirus infection of neurons in the cortex and thalamus. Viral titers from the paired right-brain hemispheres are displayed adjacent to the micrographs.

### NgR1 is not required for reovirus transmission along neural routes following intramuscular inoculation

We hypothesized that NgR1 might function to mediate reovirus spread through neural routes to the brain. To test this hypothesis, we inoculated newborn WT, JAM-A^-/-^, NgR1^-/-^, and DKO mice intramuscularly in the hind limb with 5×10^6^ PFU of T3SA-. At 1, 2, 4, and 6 dpi, mice were euthanized, and viral titers in the inoculated limb, inferior and superior spinal cord (ISC and SSC, respectively), brain, and blood were determined. Titers in the inoculated limb increased rapidly and reached significantly higher levels by 1 dpi in mice lacking JAM-A (JAM-A^-/-^ and DKO mice) compared with WT and NgR1^-/-^ mice (Fig. 4A). Titers in the limb were comparable in all genotypes by 2 dpi and remained comparable during the experimental time course (Fig. 4A). The initial rapid increase in titers in the absence of JAM-A could be due to viral entry using other receptors or increased tissue permeability in the absence of JAM-A (32), which might support more efficient infectivity. Viral titer in the blood was undetectable and reached around the detection limit by 6 to 8 dpi in all genotypes (Fig. 4B). Titers in the ISC, SSC, and brain continued to increase over the time course in all genotypes examined (Fig. 4C-E). Interestingly, titers in the ISC, SSC, and brain of mice lacking JAM-A were lower than those in WT and NgR1^-/-^ mice at multiple time points (Fig. 4C-E). These lower titers likely result from reduced hematogenous spread to neural sites in the absence of JAM-A (13,33). Importantly, titers in neural tissue of mice lacking NgR1 were comparable to those in mice expressing NgR1, suggesting that NgR1 is dispensable for reovirus neural spread.

**FIG 4.**
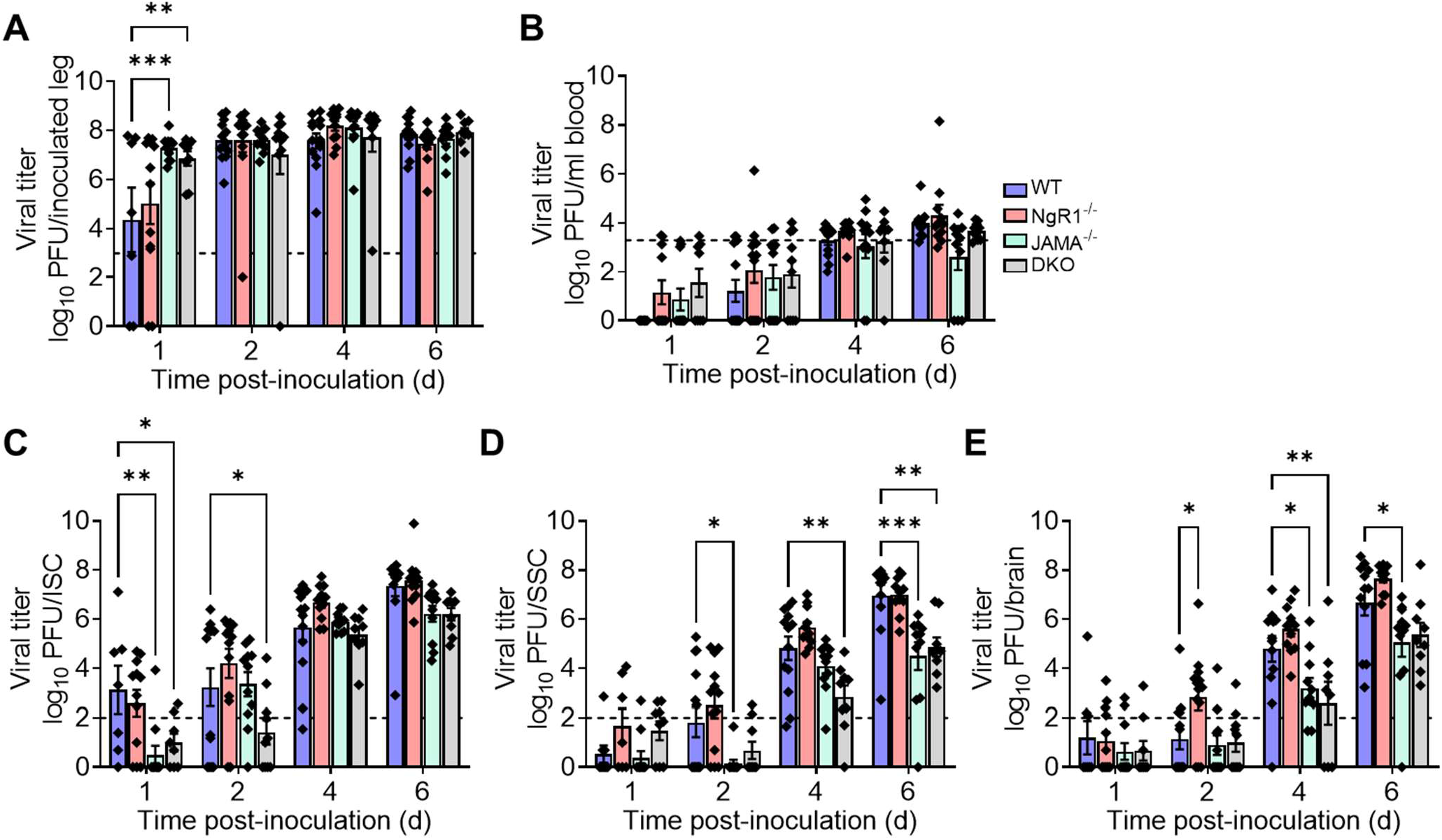
NgR1 is dispensable for reovirus neural transmission following intramuscular inoculation. WT, JAM-A^-/-^, NgR1^-/-^, and DKO mice were inoculated intramuscularly in the hindlimb with 5×10^6^ PFU of reovirus T3SA-. Titers in the (A) inoculated hindlimb, (B) blood, (C) ISC, (D) SSC, and (E) brain at the indicated intervals are shown. Dotted lines indicate the limit of detection. N = 8 to 13 animals per group for each time point. Error bars indicate SEM. Titers in tissues from knock-out animals were compared with WT at each time point. *, *P* < 0.05; **, *P* < 0.01; ***; *P* < 0.001; as determined by two-way ANOVA with Dunnett’s multiple comparisons test.

### Serotype 1 reovirus does not require NgR1 for replication in the brain

Expression of the human homolog of NgR1 in non-susceptible cells allows infection by both T1 and T3 reovirus strains (26). Therefore, we tested whether NgR1 is required for reovirus infection of the brain by T1 reovirus strain, T1L. WT and NgR1^-/-^ mice were inoculated intracranially with 100 PFU of T1L. Mice were euthanized at 3, 6, and 9 dpi, brains were removed and hemisected, and titers from the inoculated half of the brain were determined. No significant differences in virus titer were observed between WT mice and those lacking NgR1 (Fig. 5), suggesting that NgR1 is dispensable for reovirus T1L replication in the brain following intracranial inoculation.

**FIG 5.**
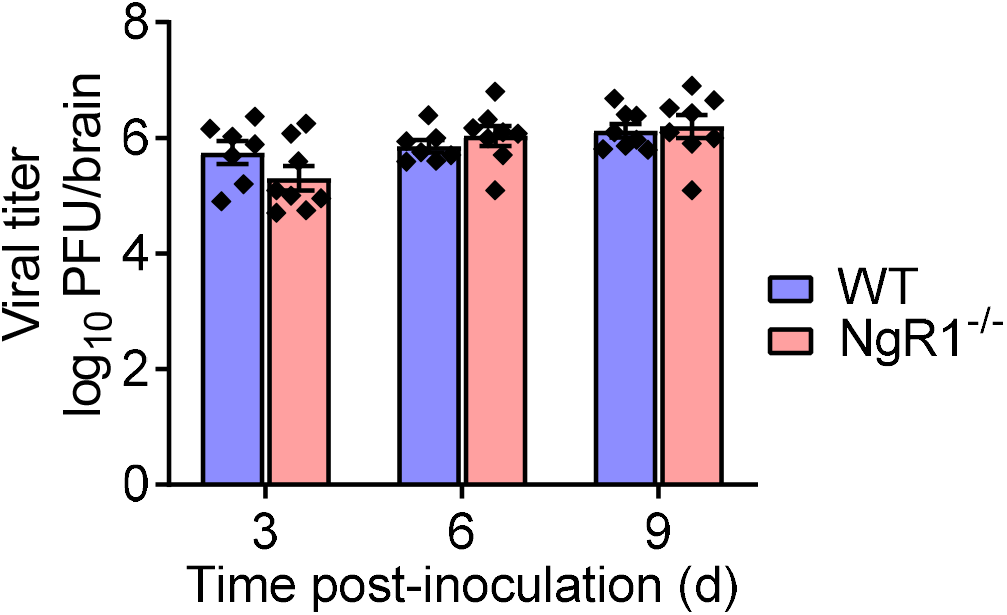
NgR1 is not required for replication of serotype 1 reovirus in the brain. Newborn WT and NgR1^-/-^ mice were inoculated intracranially with 100 PFU of reovirus T1L. At 3, 6, and 9 d after inoculation, mice were euthanized, brains were removed and hemisected, and viral titers in homogenates of the inoculated half of the brain were determined by plaque assay. Results are expressed as mean viral titers. Error bars indicate SEM. N = 7 to 8 mice per group for each time point. Differences in titer between WT and NgR1^-/-^ mice at each time point were not significant (*P* > 0.05) as determined by two-way ANOVA with Dunnett’s multiple comparisons test.

### Human but not murine NgR1 functions as a reovirus receptor

Human NgR1 directly engages reovirus virions and allows infection of non-susceptible cells (26). Given the function of human NgR1 as a reovirus receptor, the dispensability of murine NgR1 in reovirus pathogenesis was puzzling and suggests that murine NgR1 is incapable of engaging reovirus and mediating infection. We tested the capacity of CHO cells expressing murine or human NgR1 to bind and allow infection by reovirus. JAM-A was used as a positive control and the coxsackievirus and adenovirus receptor (CAR), which does not engage reovirus (24,34), was used as a negative control. A significant fraction (> 50%) of JAM-A- or human NgR1-expressing cells bound reovirus, whereas CAR- or murine NgR1-expressing cells bound reovirus poorly (Fig. 6A). Differences in reovirus binding to cells expressing CAR or mouse NgR1 were not statistically significant. The observed differences in binding are not attributable to differences in transfection or receptor-expression efficiencies, as > 75% of transfected cells expressed human or murine NgR1 on the surface, and only receptor-expressing cells were included in the determination of the percentage of reovirus-bound cells. Differences in binding paralleled differences in infectivity. JAM-A or human NgR1 expression led to infection of CHO cells, but CAR or murine NgR1 expression did not (Fig. 6B and C). These results confirm that mouse NgR1 does not engage reovirus efficiently or allow infection of non-susceptible cells.

**FIG 6.**
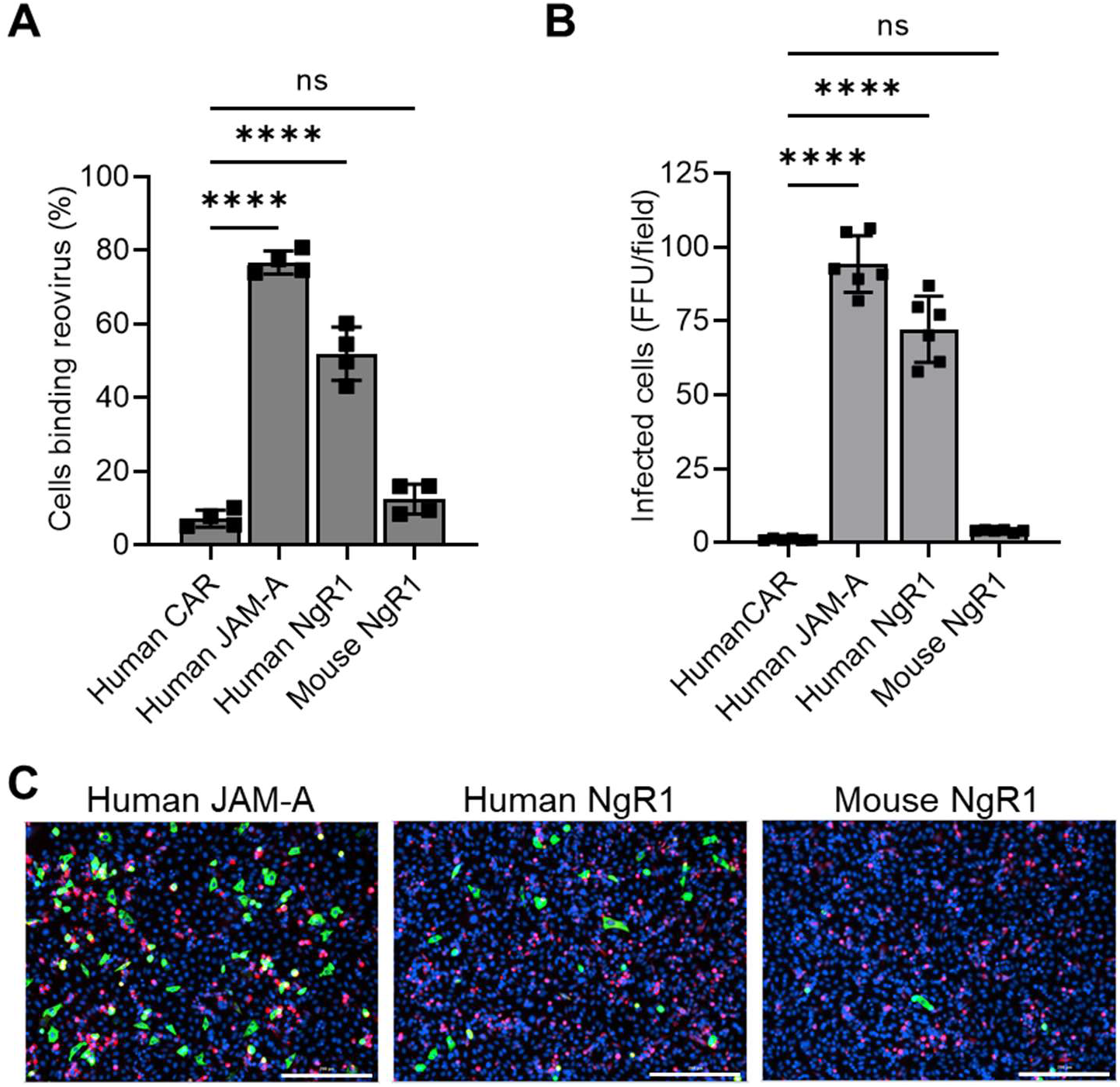
The murine homolog of NgR1 does not efficiently engage reovirus. (A) CHO cells were transfected with plasmids encoding the proteins shown and incubated with 10^5^ particles/cell of fluorescently-labeled reovirus T3SA- at 4°C for 1 h. The percentage of receptor-expressing cells bound by reovirus was quantified using flow cytometry. (B-C) Transfected CHO cells were adsorbed with 30 PFU/cell of T3SA-. (B) Infectivity was quantified at 24 h post-adsorption. Results are expressed as the mean viral values from duplicate samples of 2 independent experiments. Error bars indicate SEM. ns, *P* > 0.05; ****, *P* < 0.0001; as determined by one-way ANOVA with Dunnett’s multiple comparisons test. (C) Representative micrographs display nuclei stained with DAPI (blue), corresponding receptor expression (red), and reovirus antigen staining indicating infected cells (green). Scale bars, 200 μm.

## DISCUSSION

Serotype-specific receptor engagement dictates reovirus tropism for distinct CNS regions (15,16,18). However, the identity of receptors mediating reovirus neurotropism is unknown. Human NgR1 functions as a receptor for reovirus *in vitro* and is expressed primarily in neural tissues, suggesting that NgR1 mediates reovirus neurotropism. We tested this hypothesis by examining reovirus dissemination and replication in NgR1^-/-^ mice. The absence of NgR1 did not alter replication of neurotropic reovirus strain T3SA- in the intestine or its dissemination to the brain following peroral inoculation. Following intramuscular inoculation, which provides access to neural routes, T3SA-titers increased in the spinal cord and brain of WT and NgR1^-/-^ mice comparably, suggesting that neural dissemination routes are intact in NgR1^-/-^ mice. Viral titers in DKO mouse tissue homogenates were lower than those in WT mice and comparable to those in JAM-A^-/-^ mice, indicating that the differences in viral titers are attributable to the lack of JAM-A expression and concomitant diminished hematogenous viral spread (13,33). Reovirus T3SA-replication, tissue tropism, and virulence in the CNS were comparable in WT and NgR1^-/-^ mice, excluding a function for murine NgR1 in reovirus neuropathogenesis. These puzzling results were reconciled by our finding that human but not murine NgR1 efficiently binds reovirus and functions as a receptor *in vitro*.

The poor capacity of murine NgR1 to bind or allow reovirus infection of nonsusceptible CHO cells is in contrast with its requirement for infection of murine cortical neuron cultures reported previously (26). Receptor blockade with NgR1-specific antibodies or absence of NgR1 expression diminishes reovirus infection of primary murine cortical neurons in culture (26). Our findings recapitulate the diminished infectivity observed for T3 reovirus in cortical neurons derived from embryonic NgR1^-/-^ mice relative to those from WT mice (Fig. 2C), although the magnitude of the infectivity difference was not as striking as previously observed (26). These differences could be due to subtle variations in the culture conditions that alter expression of NgR1 or an alternative neural receptor for reovirus that allows infection in the absence of NgR1.

Our results demonstrating that murine NgR1 is dispensable for reovirus neuropathogenesis raise two important questions. First, how does murine NgR1, despite its incapacity to efficiently bind reovirus, promote infection of cortical neurons in culture? Second, what is the basis for the NgR1 requirement for infection of cortical neurons in culture but not for *in vivo* infection? If the binding of reovirus to murine NgR1 is of low affinity, then it is possible that binding to murine NgR1-expressing CHO cells at steady state may not be detectable by flow cytometry. However, neurons may express additional factors that contribute to higher-affinity binding and allow infection by reovirus. In support of this idea, while human NgR1 expression allows infection by both T1 and T3 reovirus (26), neuronal cultures allow infection only by T3 reovirus (13,18,35), suggesting that neuronal factors other than NgR1 determine reovirus tropism both *in vitro* and *in vivo*. These alternative receptors may not be expressed efficiently in neurons isolated from embryonic mice, thereby shifting the dependency to NgR1 for infection. It also is possible that murine NgR1 acts as a receptor for reovirus in specific host tissues that we did not monitor in this study. However, our results show that murine NgR1 is dispensable for reovirus neural infection.

There is precedent for the failure of viral receptors that function in cell culture to serve as receptors for *in vivo* infection. For example, Axl, Mertk, and Tyro3 function as Zika virus entry receptors and allow infection of nonsusceptible cells *in vitro* but are dispensable for Zika virus infection and tropism in mice (36). CD46 is ubiquitously expressed by nucleated cells and acts as an entry receptor for measles virus vaccine strains *in vitro* (37). However, tropism of measles virus vaccine strains in macaques is limited to specific tissues and does not correlate with CD46 expression (38). Therefore, in some cases, *in vitro* viral receptors are neither necessary nor sufficient for *in vivo* infection.

Species-specific polymorphisms in viral receptors may alter the capacity of a virus to engage receptors and establish infection in the host. Human and mouse NgR1 share structural and functional homology. Both are crescent-shaped, leucine-rich-repeat proteins that engage ligands in the CNS using conserved residues on the concave surface (27,39). However, regions and residues in human NgR1 that mediate reovirus binding are unknown. Differences in key contact residues may alter the capacity of reovirus to engage murine NgR1 while allowing human NgR1 to function as a receptor. There are many examples of engagement of host-specific receptors by viruses. Cedar virus uses murine but not human ephrin-A1 as an entry receptor, and the specificity is determined by a single contact residue in the receptor-ligand interface (40). Similarly, while the murine homolog of the hepatitis B virus receptor does not support hepatitis B virus entry, a chimeric murine receptor with a four-amino-acid replacement from the human receptor is sufficient to allow virus entry and replication (5,41). Further structural insights are required to understand differences in reovirus interactions with murine and human NgR1.

Our findings clarify functions for known reovirus receptors in pathogenesis and open doors for identification of alternative receptors that dictate CNS tropism. While JAM-A is required for establishment of viremia following reovirus replication in the intestine (13,33), both JAM-A and NgR1 are dispensable for neural spread and infection. Thus, a reovirus neural receptor that dictates tropism and neuropathogenesis remains to be identified. Importantly, our results do not exclude a function for NgR1 in reovirus infections of humans. While reovirus does not cause severe acute disease in humans, reovirus occasionally infects the CNS in children (9,10) and is implicated in the loss of immunological tolerance to dietary gluten and development of celiac disease (42). Receptors mediating reovirus infection at any site in humans are unknown, and engagement of species-specific receptors merits evaluation in this context. Reovirus has been used in clinical trials as an oncolytic agent (43), and defining the human receptors bound and their functions at distinct sites, including NgR1, is essential to precisely target oncolytic reovirus vectors. Although virus-receptor interactions often are described as specific, akin to a lock-and-key mechanism, some viruses use multiple keys to open several locks. Understanding tissue- and host-specific factors required by viruses for cell entry will inform development of specific interventions, engineering of effective vaccines, and use of viruses for targeted gene delivery.

## MATERIALS AND METHODS

### Cells and Viruses

L929 cells were grown in either suspension or monolayers in Joklik’s modified Eagle’s minimal essential medium (US Biological) supplemented to contain 5% fetal bovine serum (FBS; VWR, 97068-085). CHO cells were grown in Ham’s F12 medium (Gibco) supplemented to contain 10% FBS. Both L929 and CHO media were supplemented to contain 2 mM L-glutamine, 100 units/ml of penicillin, 100 μg/ml of streptomycin, and 0.25 μg/ml of amphotericin B. Primary cortical neurons were isolated and cultivated in parallel from brains of embryonic day 15.5 WT or NgR1^-/-^ mice as described (18). Neurons were cultivated for 5-7 days *in vitro* before inoculation with reovirus.

Reovirus strains, T1L, T3SA+, and T3SA- were prepared from laboratory stocks by plaque purification followed by passaging in L929 cells. Cesium chloride density gradients were used to purify virions from infected L929 cell lysates as described (44). Viral titers were determined by plaque assay using L929 cells (45). Purified virion particle concentrations were determined from the optical density at 260 nm (1 OD_260_ = 2.1×10^12^ particles/ml) (46).

For fluorescent labeling, reovirus particles were diluted to 6×10^12^ particles/ml in freshly prepared 50 mM sodium bicarbonate buffer and incubated with 20 μM Alexa Fluor 647 succinimidyl ester dye (Invitrogen) at room temperature for 90 min. Unreacted dye was removed by dialysis against phosphate-buffered saline (PBS) overnight at 4°C.

### Mice

C57BL/6J (WT) mice were obtained from Jackson Laboratory. JAM-A^-/-^ mice (47) provided by Dr. Thomas Sato (Cornell University, NY) and NgR1^-/-^ (48) mice provided by Dr. Stephen Strittmatter (Yale University, CT) were backcrossed with C57BL/6J background strain. JAM-A^-/-^ and NgR1^-/-^ mice were interbred to produce DKO mice. Disruption of the JAM-A- and NgR1-encoding genes was confirmed by PCR.

### Inoculation of mice

Newborn mice (2- to 3-day-old) weighing 1.4-2.3 g were inoculated with purified reovirus suspended in PBS. Mice were inoculated intracranially in the right cerebral hemisphere (18), intramuscularly in the right hindlimb, or perorally (13). Viral titers in the inocula were confirmed using plaque assays. For analyses of virulence, inoculated mice were monitored daily for weight loss and disease symptoms for 21 days and euthanized when moribund. Criteria for moribundity included immobility, seizures, paralysis, or 25% body weight loss. For determination of viral titers, mice were euthanized at various intervals post-inoculation, and organs were harvested into 1 ml of PBS and stored at −80°C. Samples were frozen and thawed twice and homogenized using a TissueLyser LT (Qiagen) instrument prior to quantification of viral titers by plaque assay using L929 cells. All animal work was conducted in accordance with Public Health Service policy and approved by the University of Pittsburgh Institutional Animal Care and Use Committee.

### Histology

Mice were euthanized at various intervals following intracranial inoculation, and brains were removed and hemisected longitudinally. The right (inoculated) brain hemispheres were used to determine viral titers and the left hemispheres were fixed using 10% neutral buffered formalin for at least 24 h, imbedded in paraffin wax, and cut into 5-μm-thick sections. Tissue sections were stained with reovirus polyclonal antiserum to visualize reovirus antigen as described (18).

### Plasmid transfection and reovirus binding

Plasmids encoding human JAM-A, CAR, and NgR1 have been described (24,26). Murine NgR1 cDNA provided by Dr. Stephen Strittmatter was sub-cloned into the pcDNA3.1+ expression plasmid using sticky-end mutagenesis and custom primers. CHO cells (5×10^4^ seeded per sample) were transfected with 1 μg of receptor-encoding plasmid using TransIT-LT1 transfection reagent (Mirusbio). Cells were used to assess reovirus binding or infection 48 h post-transfection.

For assessment of reovirus binding, transfected CHO cells were detached from cell-culture dishes using Cellstripper (Cellgro) treatment at 37°C for 15 min, quenched with culture medium, and washed once with ice-cold PBS. Cells in suspension were adsorbed with 2×10^10^ reovirus virions (Alexa 647-conjugated) at 4°C for 1 h. Cells were washed twice with FACS buffer (PBS containing 2% FBS) to remove unbound virions, and cell-bound virus was fixed using 1% paraformaldehyde. Cells were incubated for 20 min on ice and washed twice using 10 mM glycine containing PBS to quench the fixative. Cells were further incubated with the following primary antibodies to determine cell-surface expression of receptors: J10.4, mouse monoclonal antibody against JAM-A (provided by Charles Parkos, Emory University) (49); rabbit monoclonal antibody against CAR (Sinobiological, 10799-R271); goat polyclonal antibody against human NgR1 (R&D Systems, AF1208); and goat polyclonal antibody against mouse NgR1 (R&D Systems, AF1440). Secondary antibodies conjugated to Alexa Flour 488 were used to detect receptor expression. Cells were analyzed using a LSRII flow cytometer (BD Bioscience), and intensity of reovirus bound to receptor-expressing cells was quantified using FlowJo software.

### Reovirus infectivity assay

CHO cells transfected with receptor-encoding plasmids were adsorbed with reovirus T3SA-virions at a multiplicity of infection (MOI) of 30 PFU/cell, and cortical neurons in culture were adsorbed with reovirus T3SA+ at an MOI of 500 PFU/cell. Virions were diluted in PBS and adsorbed to cells at 37°C for 1 h. The inoculum was removed, cells were fixed after an additional 24-h incubation, and reovirus antigen was visualized by indirect immunofluorescence using polyclonal reovirus antiserum (50) and a fluorophore-conjugated secondary antibody. DAPI was used to stain nuclei. Cells were imaged using a LionHeart FX imager. Reovirus antigen-positive cells (fluorescence focus units; FFU) were enumerated manually for neurons and using Gen5 software for CHO cells.

## ACKNOWLEDGEMENTS

We are grateful to members of the Dermody lab for insightful discussions. Tissues were processed for histology and immunohistochemistry at the histology core at the Rangos Research Center at UPMC Children’s Hospital of Pittsburgh. Histological slides were imaged by the University of Pittsburgh Biospecimen Core.

This work was supported by U. S. Public Health Service award R01 AI038296 (T.S.D.). Additional support was provided by UPMC Children’s Hospital of Pittsburgh (P.A.) and the Heinz Endowments (T.S.D.). The funders had no role in study design, data collection and analysis, decision to publish, or preparation of the manuscript.

